# Challenges in estimating the impact of vaccination with sparse data

**DOI:** 10.1101/302224

**Authors:** Kayoko Shioda, Cynthia Schuck-Paim, Robert J. Taylor, Roger Lustig, Lone Simonsen, Joshua L. Warren, Daniel M. Weinberger

## Abstract

**Background:** The synthetic control (SC) model is a powerful tool to quantify the population-level impact of vaccines, because it can adjust for trends unrelated to vaccination using a composite of control diseases. Because vaccine impact studies are often conducted using smaller subnational datasets, we evaluated the performance of SC models with sparse time series data. To obtain more robust estimates of vaccine effects from noisy time series, we proposed a possible alternative approach, “STL+PCA” method (seasonal-trend decomposition plus principal component analysis), which first extracts smoothed trends from the control time series and uses them to adjust the outcome.

**Methods:** Using both the SC and STL+PCA models, we estimated the impact of 10-valent pneumococcal conjugate vaccine (PCV10) on pneumonia hospitalizations among cases <12 months and 80+ years of age during 2004-2014 at the subnational level in Brazil. The performance of these models was also compared using simulation analyses.

**Results:** The SC model was able to adjust for trends unrelated to PCV10 in larger states but not in smaller states. The simulation analysis confirmed that the SC model failed to select an appropriate set of control diseases when the time series were sparse and noisy, thereby generating biased estimates of the impact of vaccination when secular trends were present. The STL+PCA approach decreased bias in the estimates for smaller populations.

**Conclusions:** Estimates from the SC model might be biased when data are sparse. The STL+PCA model provides more accurate evaluations of vaccine impact in smaller populations.

## INTRODUCTION

Evaluating vaccination programs is essential to understand their benefit and to guide appropriate allocations of healthcare resources. However, it is challenging to quantify the reduction in disease rates caused by a vaccine at the population-level because various unrelated factors also affect the outcomes of interest. For example, improvements in living conditions and access to preventive care may decrease the incidence of a disease and exaggerate the effect of a vaccine. In contrast, improvements in the capacity of inpatient care services and better disease surveillance may increase the number of hospitalizations, and thus mask true vaccine-associated declines.^1^

Several methods have been proposed to control for such unrelated trends when assessing the impact of an intervention. Most commonly, linear trends (e.g., interrupted time series analysis,^2^ Holt Winter method^3^) are used to adjust for unrelated changes. A limitation of these approaches is that they assume that the pre-intervention trends are linear and continue as such in the post-vaccine period. Alternatively, a time series of a control disease can be used in a regression model to adjust for trends that influence both the disease of interest and the control disease. This approach has the advantage of capturing irregular and unexpected trends, and it draws on information about the control disease from the post-intervention period. This approach can be used to isolate the effect of the intervention if the control disease is not influenced by the intervention, if the relationship between the control disease and the disease of interest is consistent over time, and if the control captures the relevant trends. The synthetic control (SC) framework builds on this concept but combines several control diseases into a single composite and has been used to assess the impact of interventions in economics, political science, and website analytics.^4–6^ We have previously demonstrated the utility of the SC approach in estimating changes in pneumonia hospitalizations associated with the introduction of pneumococcal conjugate vaccines.^7^

When evaluating the effects of interventions using datasets with large numbers of cases, the time series of the disease of interest and the control diseases are measured with relatively little noise. With these types of data, the SC approach can effectively adjust for unmeasured confounding. In practice, however, interventions often need to be evaluated using noisy data with few cases. It is not clear whether the SC approach can successfully select an optimal set of controls and adjust for shared underlying trends when using sparser time series data.

We therefore set out to evaluate the performance of the SC approach in such settings.^6,8^ We first evaluated the ability of the SC method to quantify the impact of a 10-valent pneumococcal conjugate vaccine (PCV10) on all-cause pneumonia hospitalizations at subnational levels (e.g., regions and states) in Brazil. Based on these analyses, we found that the SC approach failed to yield reasonable counterfactual estimates when data became sparse. Therefore, we proposed an alternative approach to obtain more robust estimates of vaccine effects from sparse time series by first extracting smoothed trends from the controls. We evaluated this approach, which we called the “STL+PCA” method (seasonal-trend decomposition plus principal component analysis), using Brazil data and simulated time series data, and compared its performance with those of both the SC approach and a standard time-trend model.

## METHODS

### Hospitalization data, down-sampled data, and simulated time series data

Three types of data were used in this study: 1) national, regional, and state-level hospitalization data in Brazil, 2) down-sampled data based on the national Brazil data, and 3) simulated time series data. Detailed information on these data are provided in eAppendix 1–3, and the key information for each dataset is highlighted here.

We used a national hospital discharge database from Brazil for hospitalizations that occurred between January 2004 and December 2014.^7^ Hospitalizations were categorized using a single International Classification of Diseases (ICD) 10 code. There are 27 states in the country, which are grouped into five geographic regions: North (seven states), Northeast (nine states), Southeast (four states), South (three states), and Centre-West (four states). Children under 12 months of age and adults 80+ years of age were included in the analyses for contrast because previous studies of national-level data for Brazil demonstrated clear benefits of PCV10 in the infants but no benefit in the elderly.^7,9^ Moreover, there was a strong increasing secular trend in pneumonia hospitalizations among older adults but not among young children (eFigure 1). These different characteristics provided an opportunity to evaluate the model performance when there was no effect or a significant effect of the vaccine and with the presence of or absence of longterm trends in the time series.

To investigate how the performance of various models changes depending on the number of cases per unit time, we performed down sampling analysis.^10,11^ This approach is used to simulate how the national-level time series would behave had they been drawn from a smaller population. For example, the national population in Brazil is about 200 million, but down sampling analysis allows us to simulate a time series of a theoretical population of 20 million (i.e., down-sampling rate 10%) or 2 million (i.e., down-sampling rate 1%). Methods are described in detail in eAppendix 2. In short, we randomly subsampled the time series of ICD10 chapters from the national-level data, using the binomial distribution with the rates of 10%, 5%, 1%, and 0.25%, to simulate the population sizes of different regions and states in Brazil. This sampling process was repeated 100 times for each rate.

We also used simulated monthly time series data, which included an outcome and four control diseases, to demonstrate the performance of various models (eFigure 2). The length of time series was 120 months, and a “vaccine-associated” decline was introduced starting in month 85. There was a gradual 20% decline in the number of cases in the outcome that occurred between month 85 and the end of the time series. Both the outcome and control diseases were given an annual seasonality with different amplitude and peak timing, and the time series had a u-shaped curve that (by design) could not be captured by a standard linear trend adjustment. One of the four control diseases was a “perfect” control (blue line in eFigure 2), which means that the underlying model that generates the mean for the control is the same as the outcome, except the simulated vaccine effect is set to 0. To evaluate how the performance of the statistical models changes depending on the number of cases per unit time, we manipulated the average number of cases for the outcome at the first time point (i.e., intercept) to be roughly 8000 (100%), 800 (10%), 80 (1%), and 20 (0.25%). For each of these sample sizes, we simulated 100 time series data.

### Synthetic control (SC) model

The SC method has been described previously.^7,12^ The main outcome was the number of hospitalizations for all-cause pneumonia (ICD10 code: J12–J18). The covariates (control diseases) were disease categories based on groupings of ICD10 chapters, including both infectious and non-infectious diseases (eTable 1).^7^ Key assumptions that these control diseases need to satisfy to be valid are 1) they were not affected by PCV10 and 2) relationships between the outcome and control diseases would not change over time had PCV10 not been introduced. We therefore excluded respiratory diseases except for bronchitis/bronchiolitis (J20–J22) and disease categories that could potentially be influenced by PCV10.

The SC model was fit to monthly data from the pre-PCV10 period (2004–2009) and used to generate a counterfactual prediction for the post-PCV10 period (2010–2014). Further information can be found in eAppendix 4 while full details on the SC method are provided in Brodersen, *et al*.^12^ We used the bsts and CausalImpact packages in R version 3.4.3 (Vienne, Austria) for model fitting.^12,13^

### STL+PCA model

To address issues of sparsity in the control variables, we propose an alternative approach, “STL+PCA” model, where we first extract a long-term trend for each control variable, obtain a “composite” of these trends, and use this composite trend to adjust pneumonia rates (eFigure 3). The first step of the STL+PCA method is to extract smoothed trends from the time series of each of the control diseases using the seasonal-trend decomposition procedure based on locally weighted scatterplot smoothing (STL method).^14^ The span of the locally weighted scatterplot window can be adjusted to control the smoothness of extracted trends (eFigure 4 and eAppendix 5).

The second step is to obtain a “composite” trend among extracted trends for control diseases and to reduce the dimensionality of the total set of trends for control time series. To do so, we performed a principal component analysis (PCA) with extracted trends.^15–18^ The first principal component “PC1” is a linear combination of extracted trends for control diseases with maximum variance, which accounted for about 75–90% of the total variance in the extracted trends for all control diseases.

In the third and final step, we use PC1 as a covariate in a regression model for the prevaccine data and to generate the counterfactual for the post-vaccine period. We only included PC1 in the model, because PC1 explained most of the variance (75–90%) and the rest of the principal components were just capturing the remaining noise. More information can be found in eAppendix 5. We validated the STL+PCA model by performing a cross validation analysis with the pre-vaccine data (eAppendix 8).

### Time-trend model

The structure of the model is described in eAppendix 6.

### Evaluation of the impact of vaccine

Each of the models was fit to pneumonia time series from the pre-vaccine data to establish a relationship between pneumonia and the covariates. We then estimated the number of pneumonia cases that would have occurred without vaccination in the post vaccine era (i.e., counterfactual), assuming that the relationships between the outcome and covariates were consistent. We calculated the rate ratio (RR) of the observed to the counterfactual pneumonia hospitalizations during the evaluation period, which was 2013–2014 for the Brazil data (i.e., 36–60 months following the introduction of PCV10) and the last two years for the simulated time series data (i.e., 12–36 months after the simulated introduction of the vaccine). Posterior medians and 2.5^th^ and 97.5^th^ percentiles were reported as point estimates and 95% credible intervals (CIs) of RR, respectively.

## RESULTS

### Performance of the models with national, regional, and state-level data from Brazil

The time series for all-cause pneumonia hospitalizations among children under 12 months of age in Brazil showed strong seasonality with peaks occurring in the winter (eFigure 1A). All models (the SC, time-trend, and STL+PCA model) detected a significant decline in the number of pneumonia hospitalizations after the introduction of PCV10 in 2010 at the national level in this age group. National-level estimates of RR were 0.72 (95% CI: 0.67, 0.77) by the SC model, 0.85 (95% CI: 0.73, 0.99) by the time-trend model, and 0.83 (95% CI: 0.71, 0.96) by the STL+PCA model. Similarly, all models estimated significant declines in most of the regions and states in this age group (Figure 1A, C, and E).

**Figure 1.**
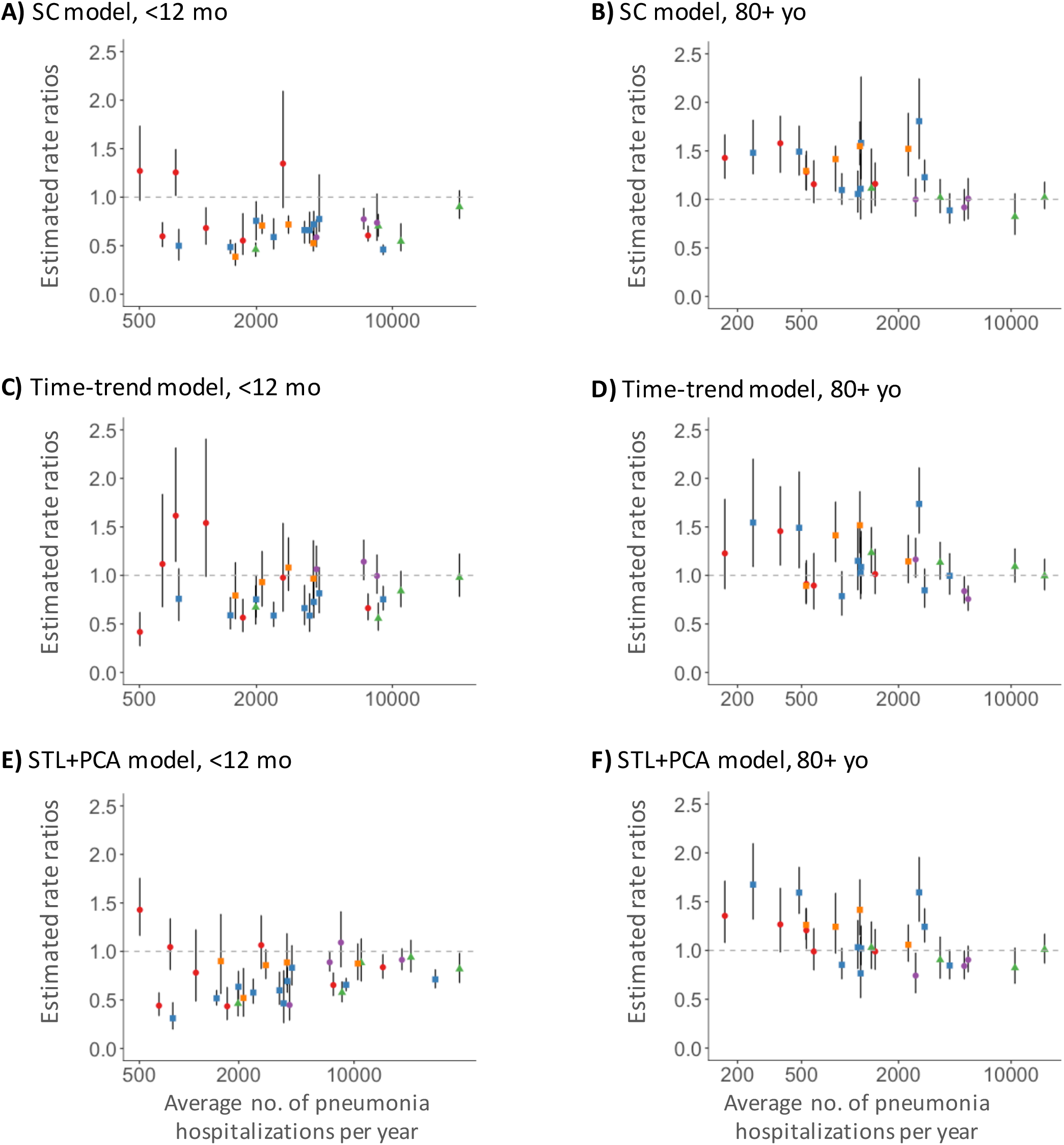
Estimates of State-level Rate Ratios by the Average Number of All-cause Pneumonia Hospitalizations in Brazil. States in the North region are represented in red, the Northeast region in blue, the Southeast region in green, the South region in purple, and the Centre-West region in orange. Abbreviations: SC, synthetic control; STL, seasonal-trend decomposition procedure based on locally weighted scatterplot smoothing; PCA, principal component analysis.

Time series data for the older age group showed a complex pattern; all-cause pneumonia hospitalizations, as well as some control diseases, began to increase several years prior to the introduction of PCV10 and continued to increase until the end of the study period (eFigure 1B). At the national level, all models adjusted for this unexplained long-term increasing trend and found no statistically significant effect of PCV10. Estimated national-level RRs were 0.95 (95% CI: 0.82, 1.12) by the SC model, 1.06 (95% CI: 0.90, 1.23) by the time-trend model, and 1.02 (95% CI: 0.88, 1.18) by the STL+PCA model. Repeating this analysis for the elderly at the state level, however, the SC model generated RRs that were significantly greater than one in 12 (46%) states, which is not epidemiologically plausible. Most of these states were found to have relatively few pneumonia hospitalizations (Figure 1B). A mean squared error (MSE, eAppendix 7) of state-level RRs compared to the national estimate was 0.148 by the SC model. State-level RRs estimated by the time-trend and STL+PCA model were less biased, resulting in smaller MSEs (0.073 for the time-trend model and 0.076 for the STL+ PCA model). The number of states with RRs greater than one decreased to 7 (28%) and 8 (32%) states by the time-trend and STL+PCA model, respectively (Figure 1D and F).

### Performance of the models with down-sampled Brazil data

To further examine the effect of smaller sample size on the accuracy of the estimates from these models while setting aside complicated factors affecting real-world data, we conducted a down sampling analysis of the national time series from Brazil. For adults age 80+ years, estimated RRs generated by the SC model rapidly diverged from the national estimate (the “ground truth”), as we sampled successively fewer cases (Figure 2A). As a result, MSE increased exponentially as the data became sparse (Figure 3).

**Figure 2.**
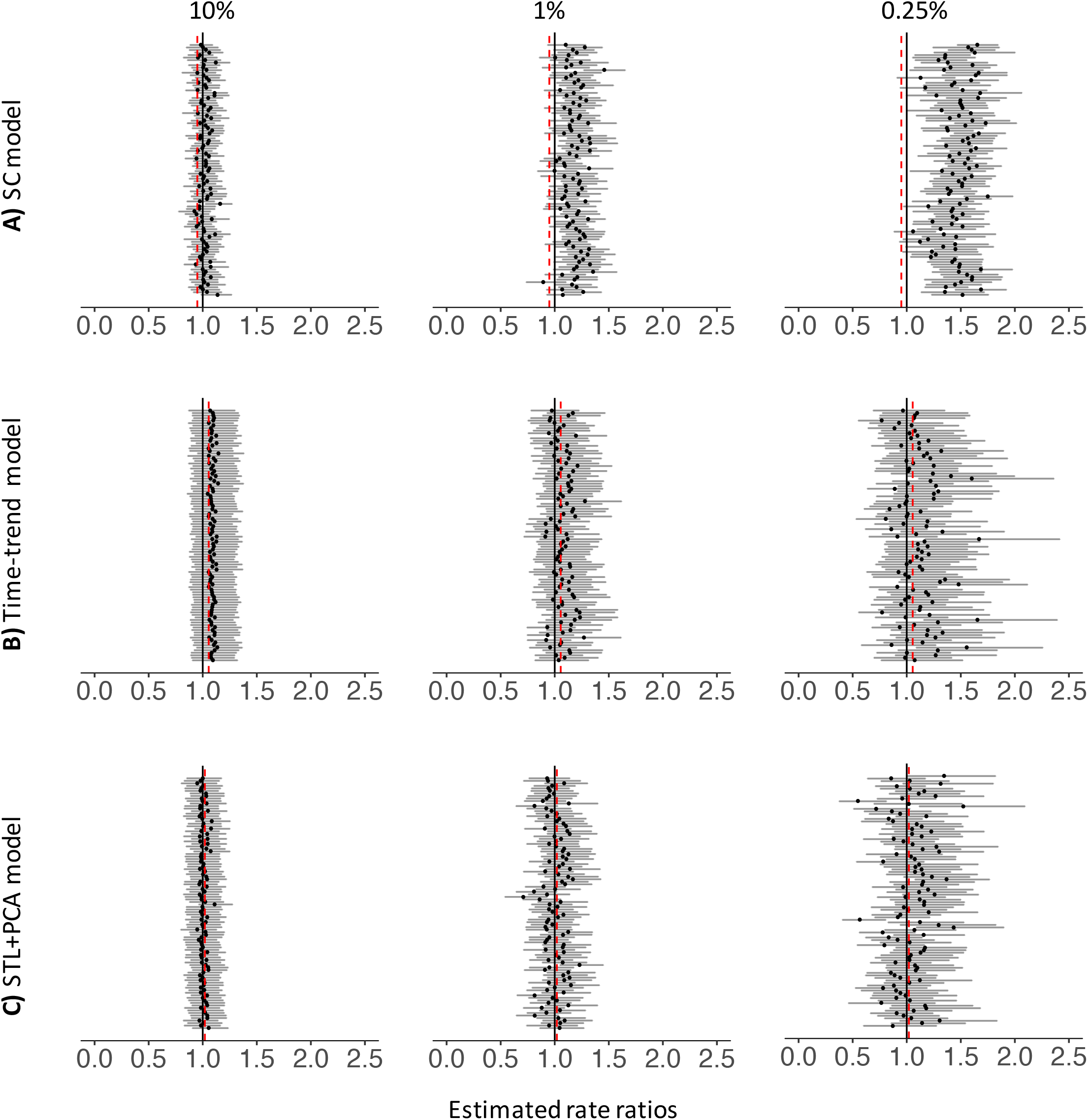
Estimated Rate Ratios for Down-sampled Datasets (80+ yo, Brazil). Each black dot represents a RR estimated for each down-sampled dataset. Dark grey bars associated with these dots represent 95% credible intervals for RRs. The percentages at the top represent the down-sampling rates. Black vertical lines represent the null value (RR=1) and red dashed lines represent national estimates of RR generated by each type of the model. Abbreviations: RR, rate ratio; SC, synthetic control; STL, seasonal-trend decomposition procedure based on locally weighted scatterplot smoothing; PCA, principal component analysis.

**Figure 3.**
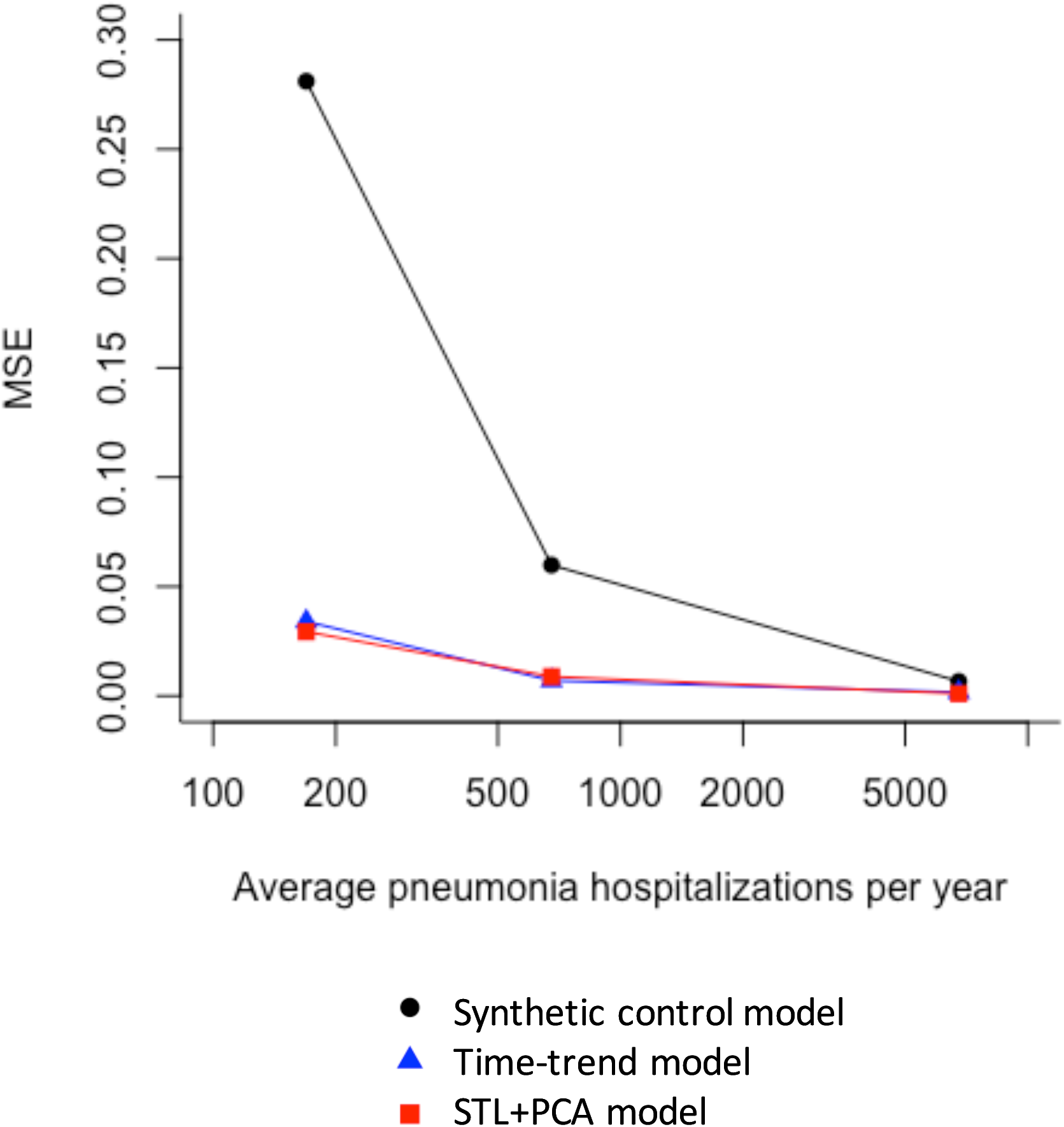
Mean Squared Errors of Estimated Rate Ratios from Down-sampled Datasets (80+ yo). Abbreviations: MSE, mean squared error; STL, seasonal-trend decomposition procedure based on locally weighted scatterplot smoothing; PCA, principal component analysis.

With the national data for the elderly, the SC model selected three control diseases on average (range: one to nine control diseases). However, when the down-sampling rate was 0.25%, the SC model did not select any control diseases in the final model in 47 of the 100 downsampled datasets (i.e., model only had an intercept and seasonal terms), only one control disease in 49 datasets, and two control diseases in the remaining four datasets. These results demonstrate that the SC model failed to identify an appropriate set of control diseases and thus failed to adjust for unmeasured confounding in the datasets down-sampled to resemble the smaller states. In contrast, the time-trend and STL+PCA method successfully adjusted for a long-term increasing trend and corrected the bias even when the data became sparse (Figure 2B and C). MSEs remained small across all down-sampling rates (Figure 3).

For children less than 12 months of age, where there was no strong secular trend, performance of all three models was not dramatically affected by sample size (eFigure 5). MSEs remained small for all models, regardless of the down-sampling rate (eFigure 6). However, the average number of control diseases selected in the SC model was zero in 88 out of 100 downsampled datasets when the down-sampling rate was 0.25%.

### Performance of the models with simulated time series data

First, we fit a model with a single predictor, the “perfect” control, which had the exact same trend as the outcome during the pre-vaccine period, and tested whether this model was able to recover the true impact of the vaccine (RR=0.8) in the simulated data. Because of this ideal, but not realistic, relationship between the outcome and the predictor, this model was expected to generate the best possible counterfactual. RRs yielded by this model tightly lined up around the true value, even when the data became sparse (Figure 4A). Next, we fit the model which included three controls, but not the perfect control, as predictors (“Unsmoothed control model” in Figure 4B and eFigure 7). This model failed to converge when the data size was large, because of a strong collinearity among those controls. As the data became sparse, estimated RRs moved away from the null and became significantly greater than the true RR (Figure 4B). The traditional time-trend model could not capture the u-shaped trend, as expected, and estimated RRs did not cover the true value regardless of sample size (Figure 4D).

**Figure 4.**
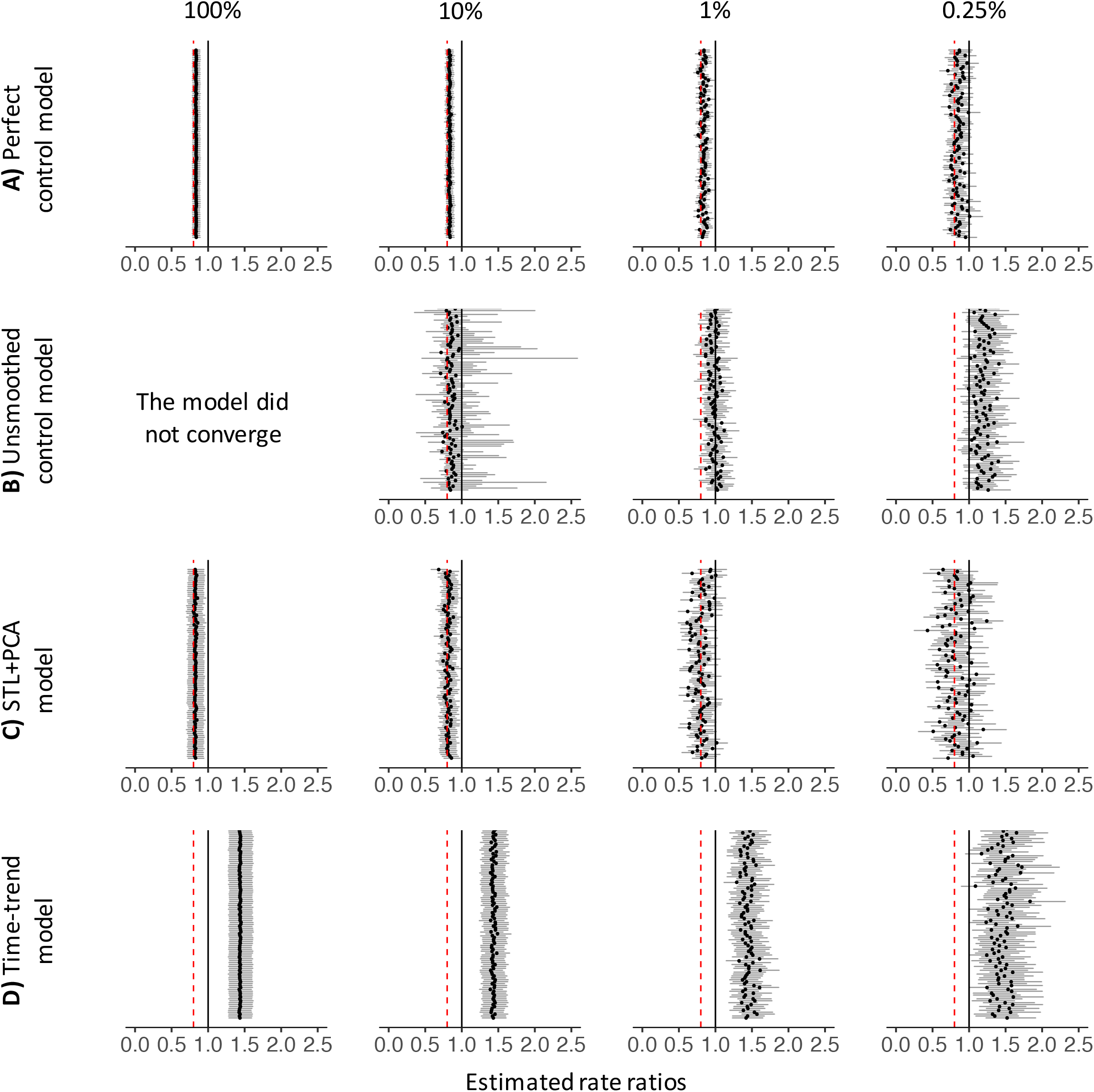
Estimated Rate Ratios for Simulated Time Series Data. Each black dot represents a RR estimated for each simulated dataset. Dark grey bars associated with these dots represent 95% credible intervals for RRs. The percentages at the top represent the sample size. Black vertical lines represent the null value (RR =1) and red dashed lines represent the true value of RR (0.8). Abbreviations: STL, seasonal-trend decomposition procedure based on locally weighted scatterplot smoothing; PCA, principal component analysis.

The STL+PCA model successfully recovered the true RR, even when the number of cases became smaller. Although point estimates of RR started to diverge gradually as data became sparse, 90% and 81% of the estimated RRs covered the true RR in their 95% CIs when the data size was 1% and 0.25%, respectively (Figure 4C). MSEs remained small regardless of the number of cases per unit time and were comparable to those for the perfect control model (eFigure 7).

## DISCUSSION

Our analyses showed that when evaluating the effects of vaccines on rates of disease using time series data, the SC model was able to successfully adjust for underlying trends in the data when the control variables had relatively little noise. However, the SC model may not be able to do so when the time series are sparse. As a result, the SC model may fail to adjust for unmeasured confounding and may generate biased estimates of the impact of an intervention. These biases tended to become stronger as sample size decreased. This was particularly a problem when there was a strong secular trend, as was observed among the elderly in Brazil. As a possible solution, we decomposed the control variables and extracted the long-term trend, and then used this long-term trend as the control. This approach led to decreased bias in the estimates of vaccine impact.

When the data did not have a long-term secular trend (i.e., among <12 month old children in our study), the sample size did not affect the performance of the SC model (eFigure 5A). It should be noted, however, that this is not because the SC model worked well in the young age group. In fact, similar to the old age group, the SC model also failed to select an appropriate set of control diseases in this age group. However, due to the lack of a secular trend, the intercept-only model was able to generate a reasonable counterfactual in this instance. We suspected that the variable selection process in the SC model did not work with the sparse data because the time series data became noisier, and many control diseases had zero cases or only a few cases per unit time in the most heavily down-sampled datasets. To test this hypothesis, we introduced the national-level control diseases in the SC model in addition to the down-sampled control diseases. This process helped the SC model to select an optimal set of control diseases, and as a result, the bias in estimated RRs was successfully corrected (eFigure 8). This analysis suggested that the problem of the variable selection may be attributed to the sparse data for control diseases but not the outcome. When data became sparse, there was more measurement error and random variation in the covariates, which biased their coefficients towards zero.^19^ As a result, the SC model failed to adjust for trends using the sparse control time series.

We proposed the “STL+PCA” model as a possible solution to the problem of data sparsity. The approach involves first extracting trends from the control time series, then using these smoothed trends to adjust pneumonia rates. Using both simulated and real-world data, we demonstrated that the “STL+PCA” approach helps to reduce the impact of sparseness and to decrease bias in the estimates for smaller populations. The first step, STL decomposition, makes it easy to identify a long-term trend in noisy time series for control diseases. The second step, PCA, allows us to find a projection that explains the maximum variability of the outcome (i.e., PC1). Users can then simply fit a regression with PC1 and generate counterfactual for the postvaccine period. Both STL and PCA are widely used and readily available in the major statistical software. Similar to the SC model, users can include all control diseases in this STL+PCA model, as long as the chosen controls satisfy the key assumptions (i.e., are not affected by the vaccine and relationships with the outcome would not have changed had the vaccine not introduced). An important disadvantage of the STL+PCA model is that it is no longer straightforward to interpret relationships between the outcome and control diseases, as the original time series for each control disease is not directly used as a covariate in the model. That is problematic if users want to understand associations between the outcome and control diseases, but is less of a concern if one’s objective is to make a robust counterfactual and quantify the impact of vaccine.

The performance of the STL+PCA model was found to be superior to those for the SC model and time-trend model, which was demonstrated by the analysis with the simulated time series data and the down sampling analysis for 80+ years-old in Brazil. With regard to the state-level data in Brazil, estimates of RR for the elderly generated by the STL+PCA model seemed to be quite similar to those generated by the time-trend model. This was because the long-term trend observed in this real-world data showed an almost perfect linear increase, which was successfully captured by both the time index variable and PC1 in those models. However, as demonstrated by the simulated time series data, the time-trend model does not work well when the long-term trend is not perfectly linear, suggesting that the STL+PCA model is more flexible than the time-trend model.

One might argue that the Bayesian variable selection process should be allowed to choose a few appropriate control disease trends to be included in the regression model, instead of performing PCA and finding the “composite” of the trends. However, the model did not converge due to the strong collinearity among extracted trends, which made it difficult to use the Bayesian variable selection. One might also argue that we should omit the STL step and perform PCA with original time series for control diseases. This approach, however, did not generate a reliable counterfactual when the original data were noisier. We found that the STL step, which allowed us to isolate long-term trends from seasonality and the remainder component was a key step to generate a robust counterfactual for sparse and noisy time series from small populations.

Another possible way to reduce the impact of sparseness is to aggregate monthly data into quarterly data and fit the SC model. This simple process increases the number of observations per unit time, thereby allowing the SC model to create an optimal composite of control diseases. This approach worked well with the Brazil data (eFigure 9). However, it may not be a good solution when the pre-vaccine data are limited. Using lower resolution time series reduces the number of data points, which makes it difficult to establish relationships between the outcome and the synthetic control. Alternative approaches could involve using a latent variable model to explicitly model the observation process, using a spatial model to borrow statistical information between adjoining localities or using model stacking approaches.^20,21^

In conclusion, the STL+PCA method could be an effective tool to infer the causal impact of vaccines and other public health interventions, and works well with sparse data. This STL+PCA model will enable us to quantify the impact of interventions more accurately, especially for small populations, and will allow us to address various public health questions using population health data that are readily available.

## FIGURE LEGENDS

**eTable 1.**
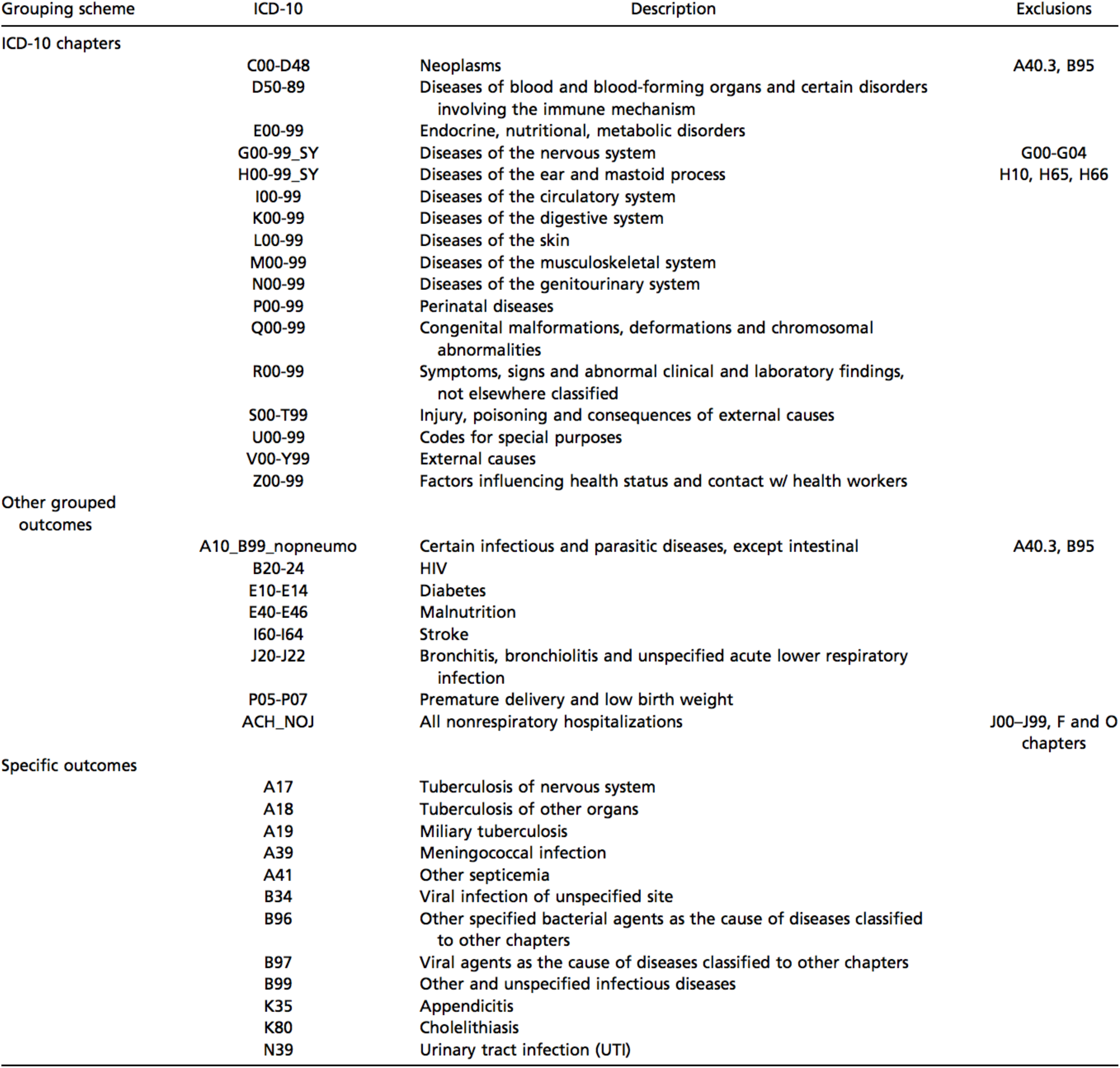
List of Control Diseases Included in the Synthetic Control Model.

**eFigure 1.**
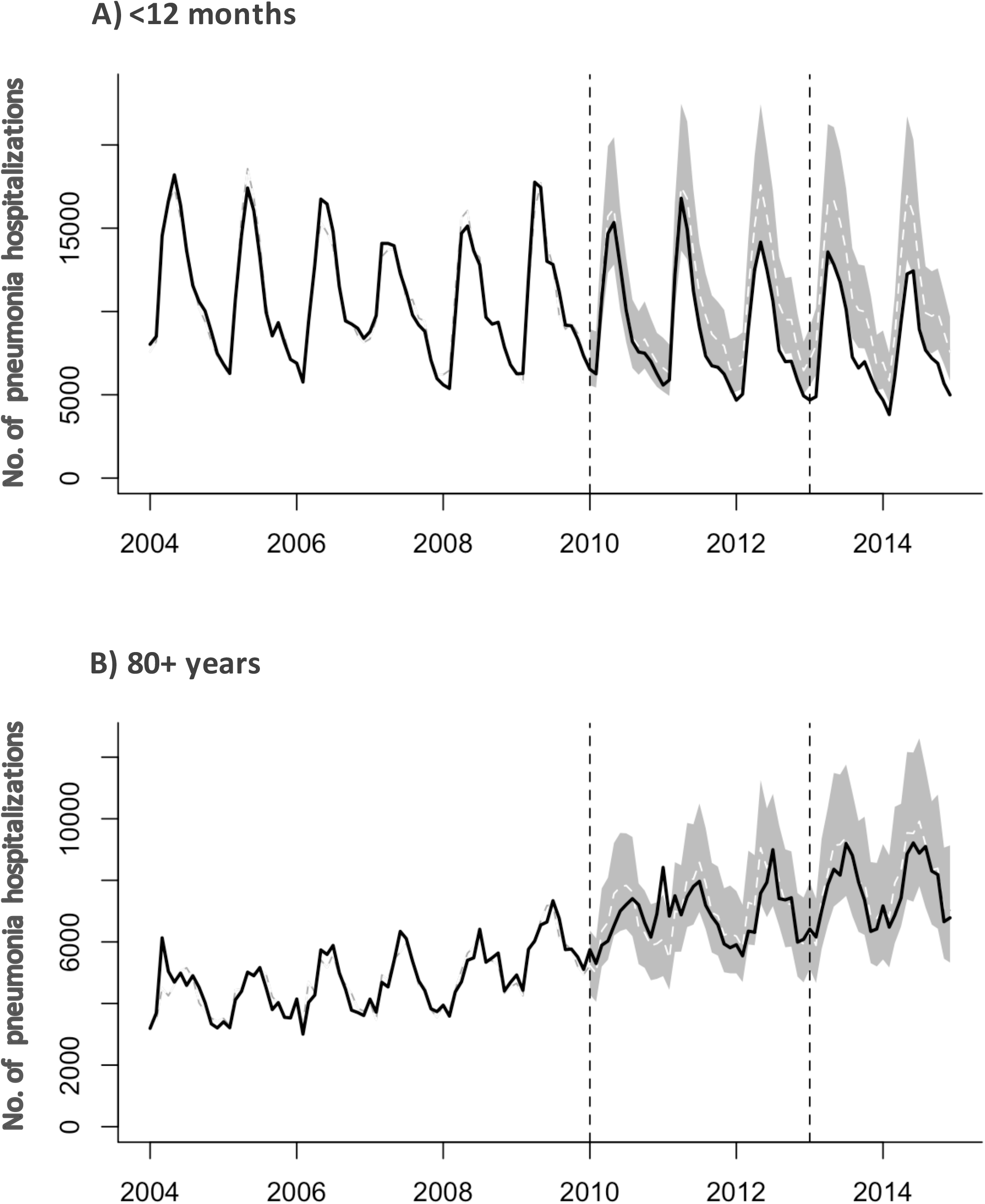
National-level Time Series of Observed and Counterfactual All-cause Pneumonia Hospitalizations Generated by the Synthetic Control Model for (A) Cases <12 Months and (B) 80+ Years of Age in Brazil. The observed number of pneumonia hospitalizations is represented by black lines. The counterfactual number of pneumonia hospitalizations and their 95% credible intervals are represented by white dashed lines and grey areas. Vertical dashed lines indicates the timing of PCV10 introduction (January 2010) and the start of the evaluation period (January 2013). Abbreviations: PCV10, 10-valent pneumococcal conjugate vaccine.

**eFigure 2.**
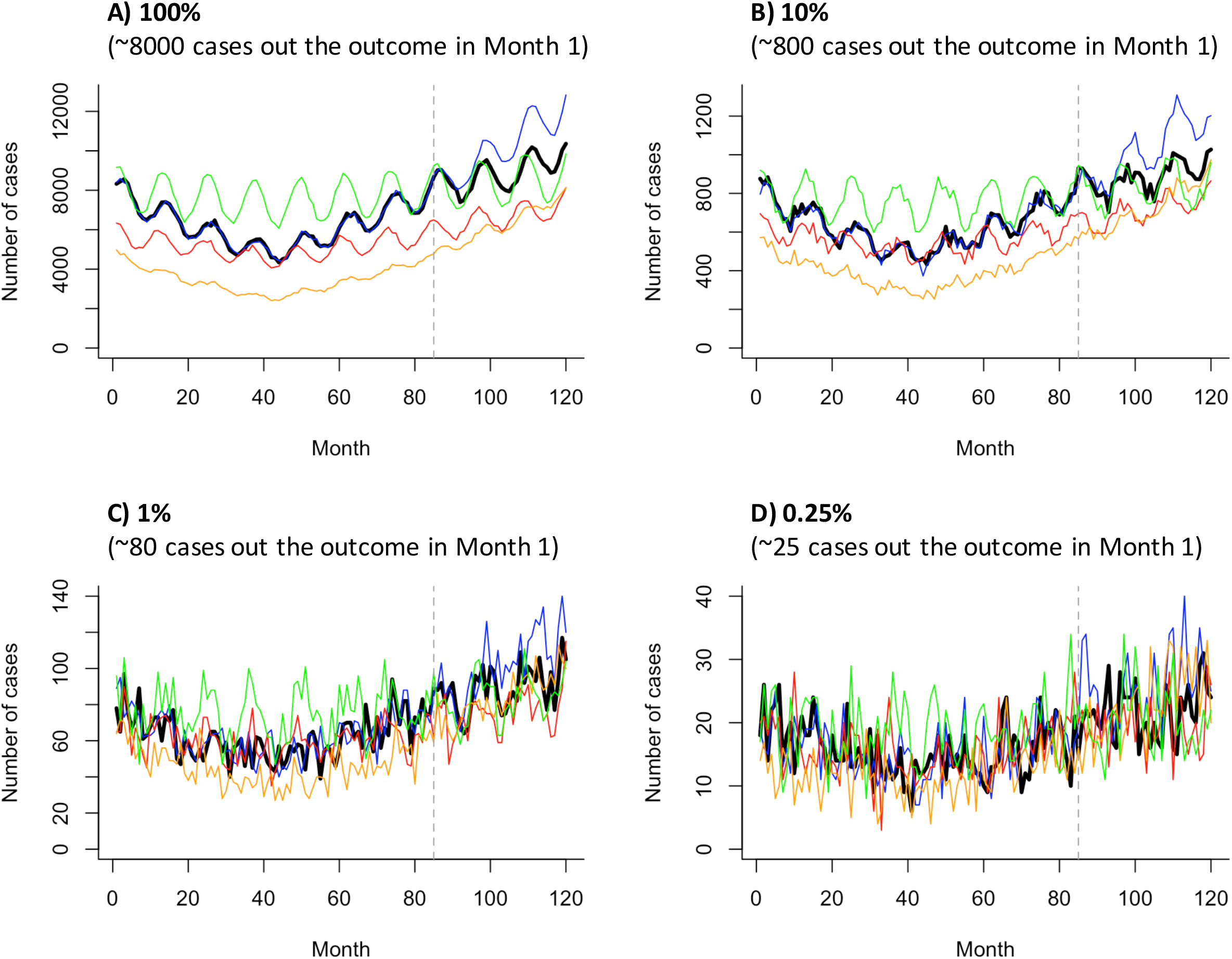
Simulated Monthly Time Series Data From a Single Simulation. The outcome is in black, the perfect control is in blue, and the remaining controls are in green, orange, and red. Percentages at the top of panels represent the sample size. Vertical dashed lines represent the timing of the simulated vaccine introduction (Month 85).

**eFigure 3.**
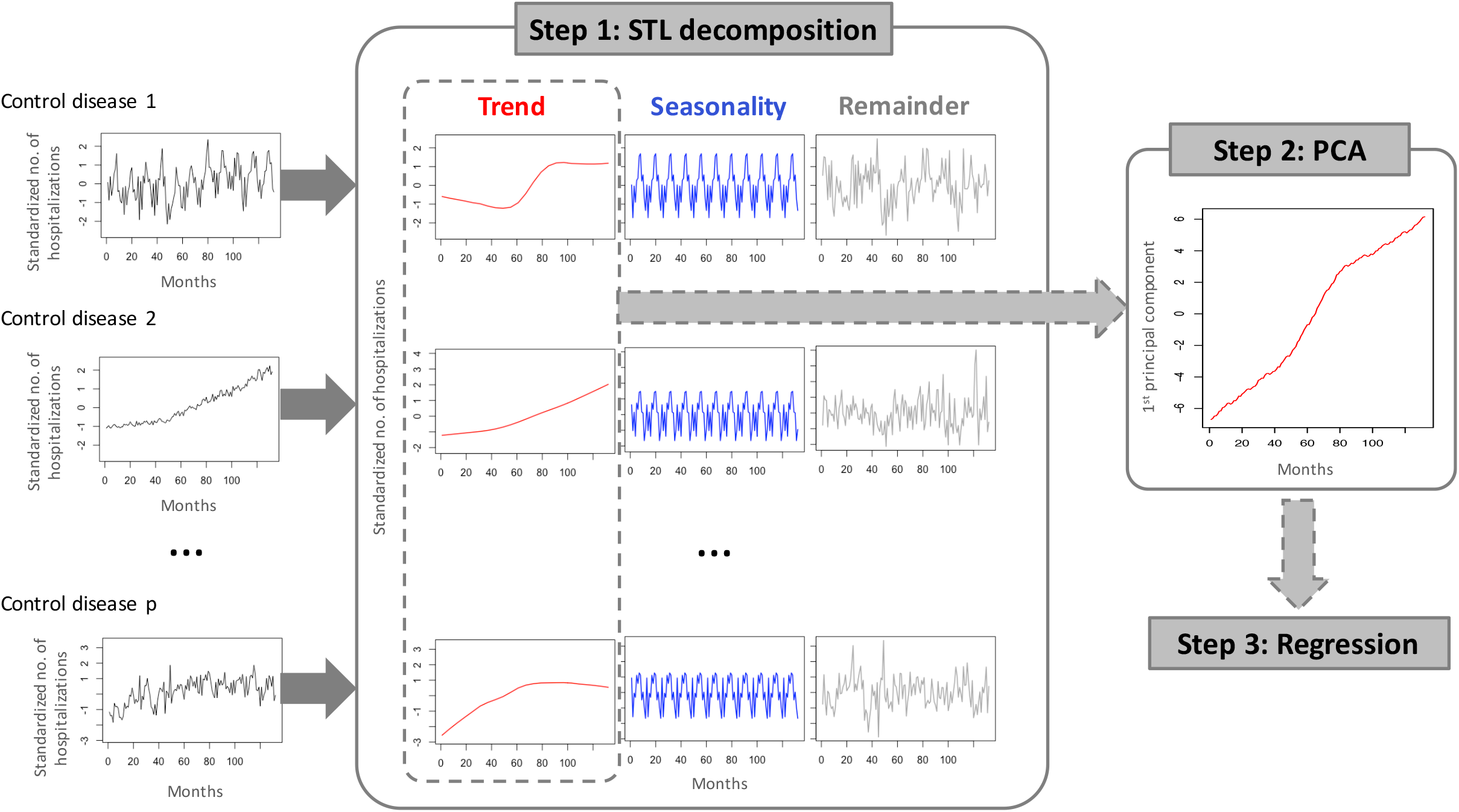
Diagram of the STL+PCA Model. Abbreviations: STL, seasonal-trend decomposition procedure based on locally weighted scatterplot smoothing; PCA, principal component analysis.

**eFigure 4.**
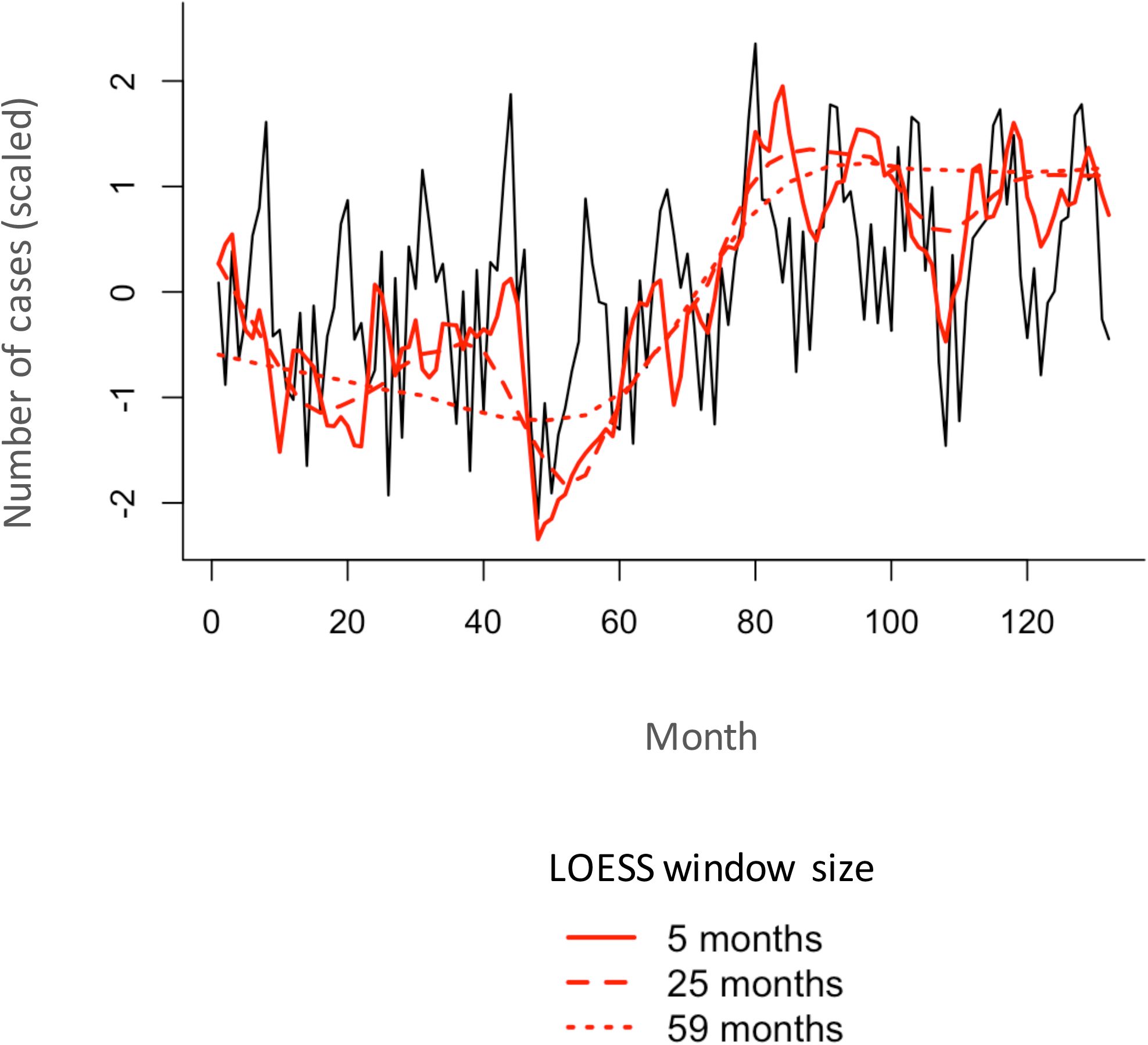
Example of STL Decomposition Using Different Sizes of the LOESS Window (ICD10 code: I00_99, 80+ yo in Brazil). Observed number of hospitalizations is in black, and trends extracted by the STL method using different sizes of the LOESS window (5, 25, and 59 months) are in red. Abbreviations: LOESS, locally weighted scatterplot smoothing; STL, seasonal-trend decomposition procedure based on locally weighted scatterplot smoothing.

**eFigure 5.**
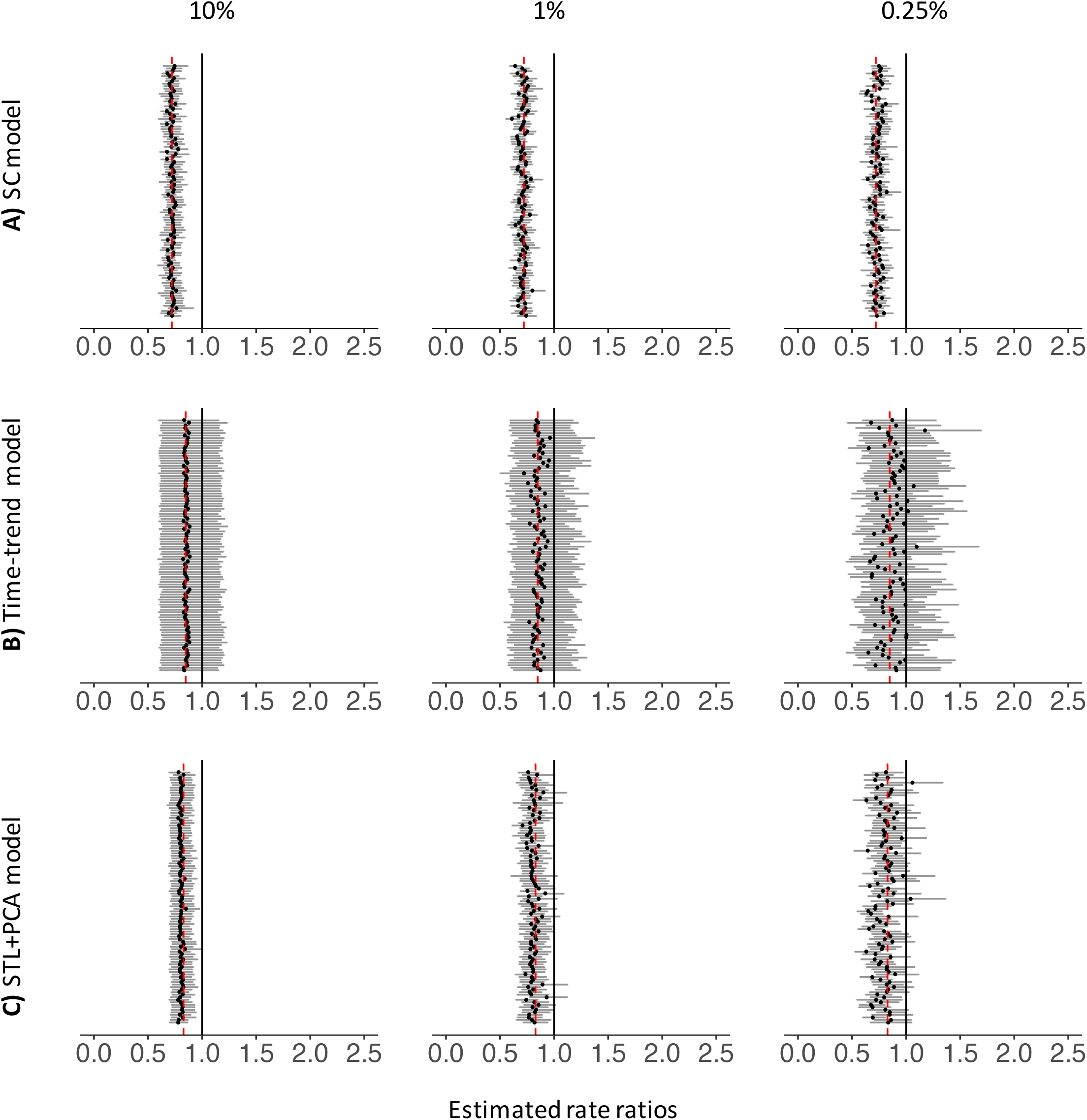
Estimated Rate Ratios for Down-sampled Datasets (<12 mo, Brazil). Each black dot represents a RR estimated for each down-sampled dataset. Dark grey bars associated with these dots represent 95% credible intervals for RRs. The percentages at the top represent the down-sampling rates. Black vertical lines represent the null value (RR=1) and red dashed lines represent national estimates of RR generated by each type of the model. Abbreviations: RR, rate ratio; SC, synthetic control; STL, seasonal-trend decomposition procedure based on locally weighted scatterplot smoothing; PCA, principal component analysis.

**eFigure 6.**
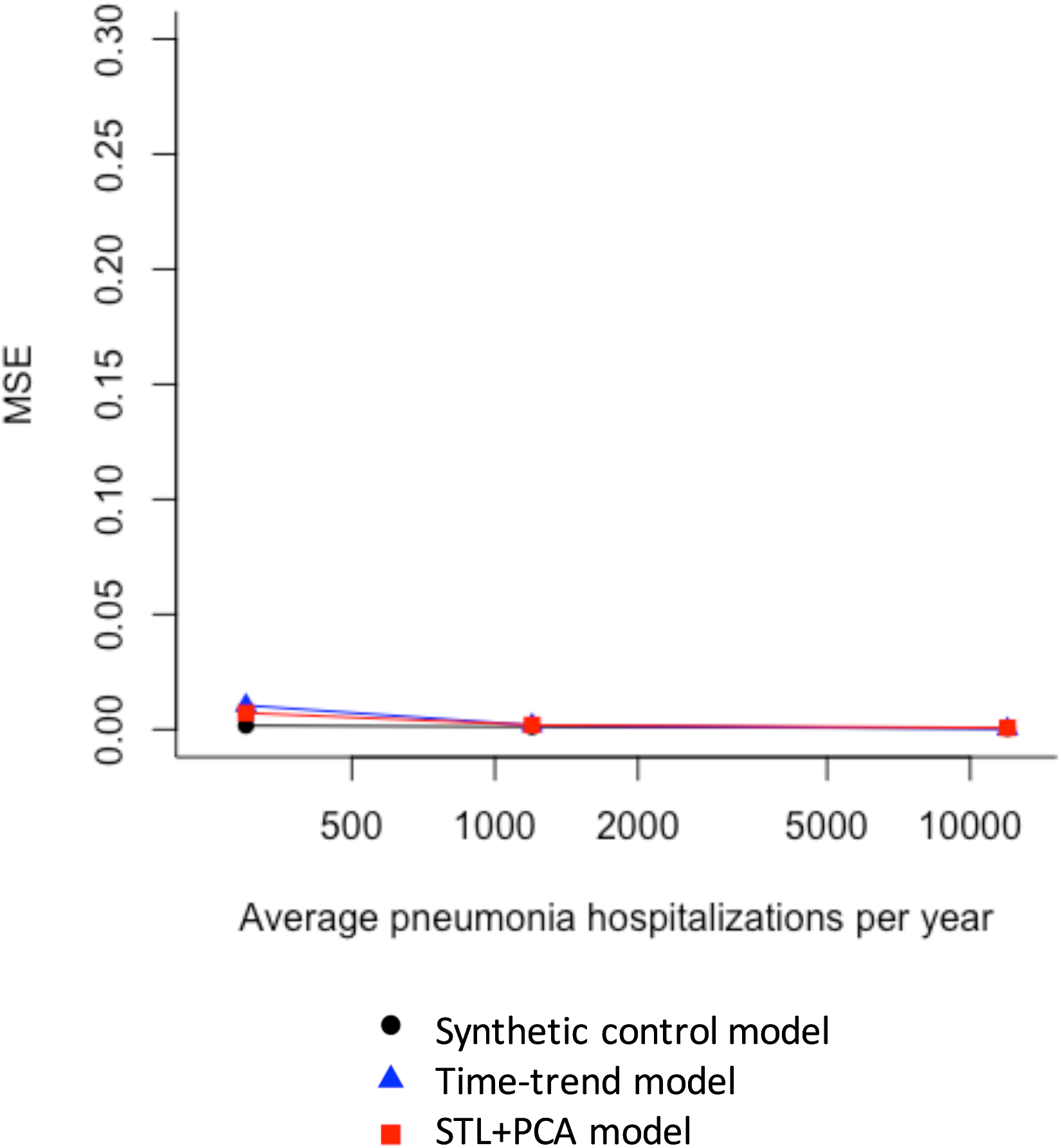
Mean Squared Errors of Estimated Rate Ratios from Down-sampled Datasets (<12 mo, Brazil). Abbreviations: MSE, mean squared error; STL, seasonal-trend decomposition procedure based on locally weighted scatterplot smoothing; PCA, principal component analysis.

**eFigure 7.**
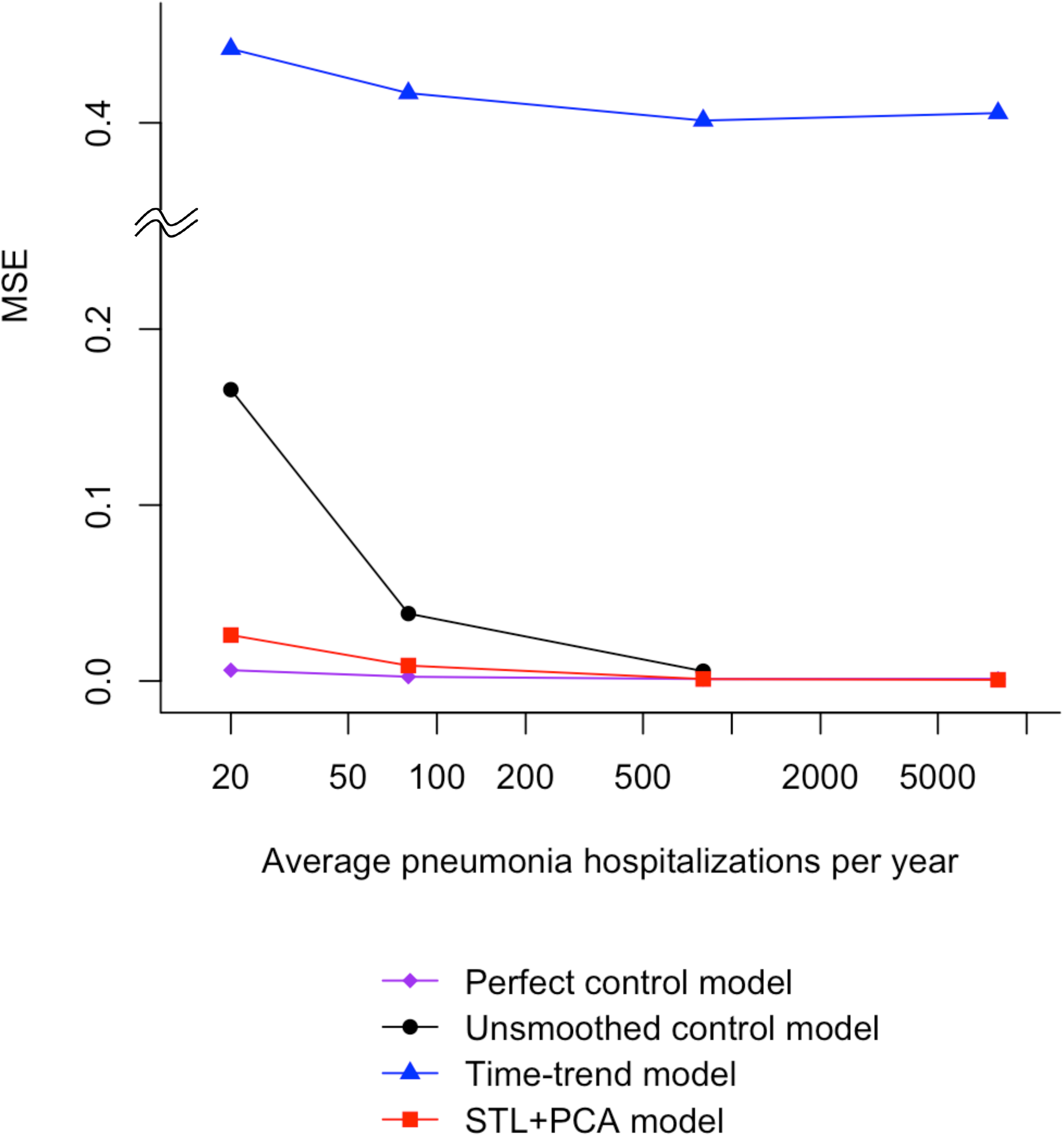
Mean Squared Errors of Estimated Rate Ratios from Simulated Time Series Data. Abbreviations: MSE, mean squared error; STL, seasonal-trend decomposition procedure based on locally weighted scatterplot smoothing; PCA, principal component analysis.

**eFigure 8.**
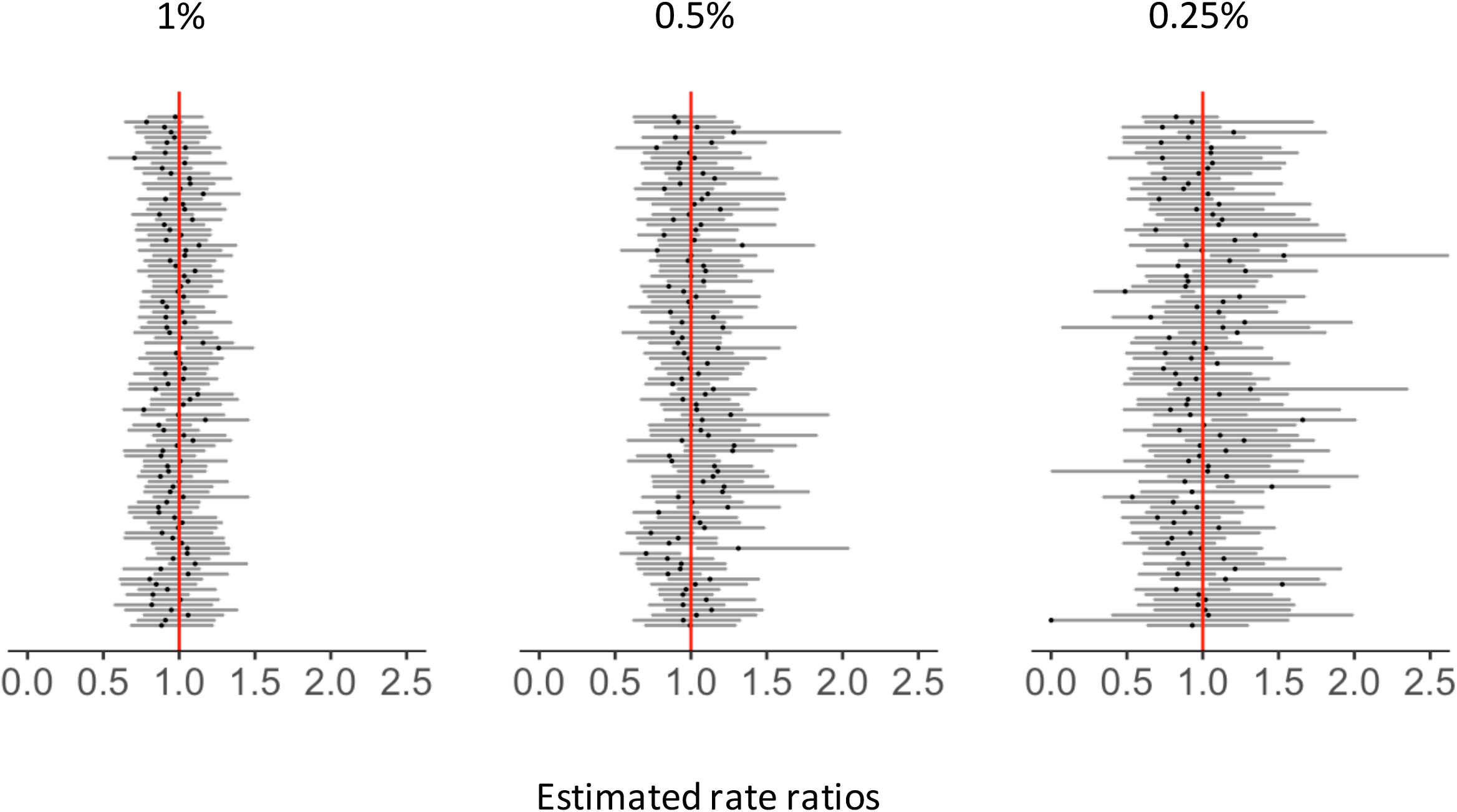
Rate Ratios for Down-sampled Datasets Estimated by the Synthetic Control Model Including Both National-level Covariates and Down-sampled Covariates (80+ yo, Brazil). Each black dot represents a RR estimated for each down-sampled dataset. Dark grey bars associated with these dots represent 95% credible intervals for RRs. Vertical red lines represent the null value (RR=1). The percentages at the top represent the down-sampling rates.

**eFigure 9.**
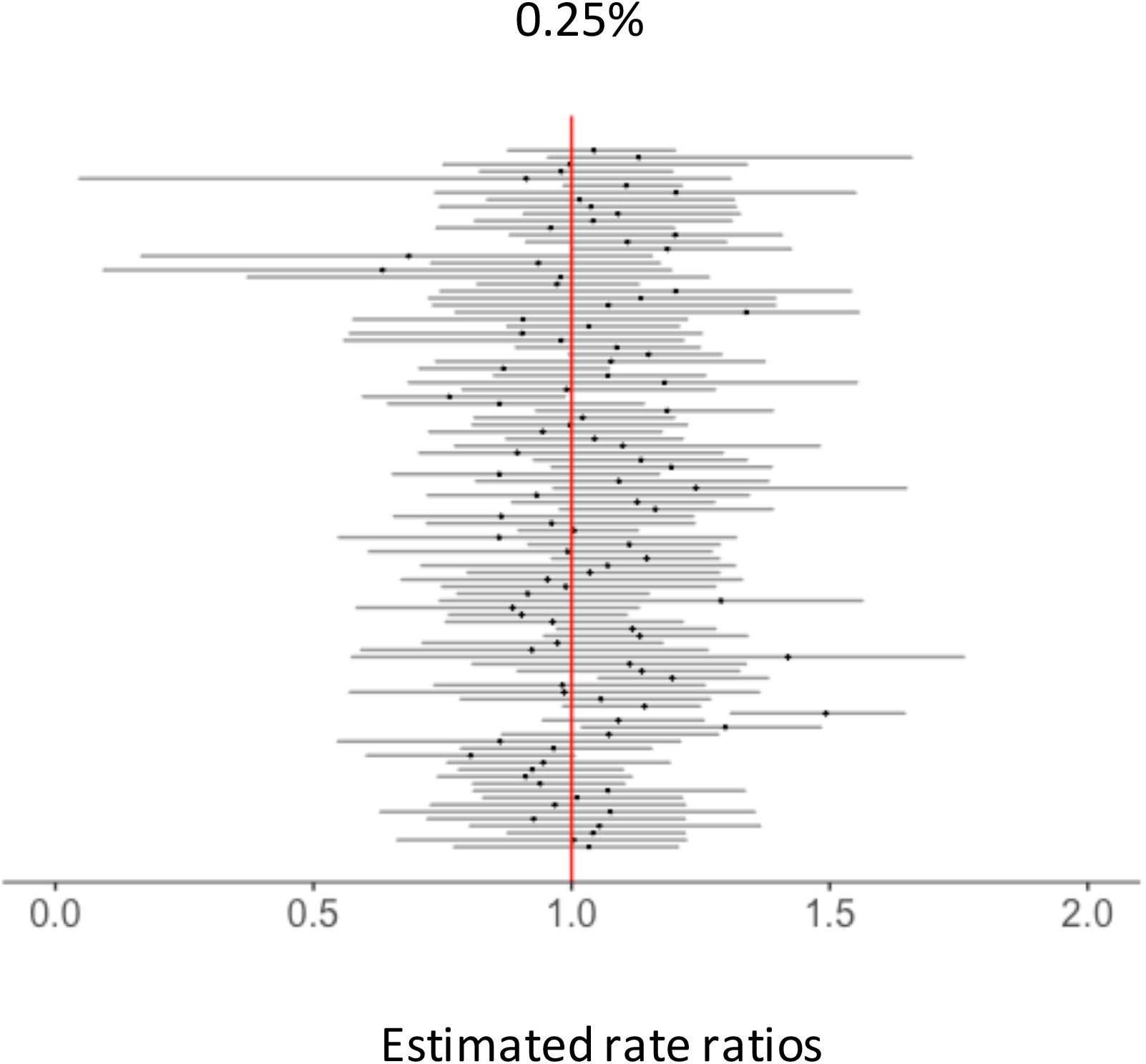
Rate Ratios for Down-sampled Datasets Estimated by the Synthetic Control Model Using Quarterly Data for 80+ yo, Brazil. Each black dot represents a RR estimated for each down-sampled dataset. Dark grey bars associated with these dots represent 95% credible intervals for RRs. Vertical red lines represent the null value (RR= 1). The percentages at the top represent the down-sampling rates.

## SUPPLEMENTAL DIGITAL CONTENT

### eAppendix 1: Additional information on hospitalization data in Brazil

The national hospital discharge database covers the majority of the national population in Brazil (e.g., 82% in 2012).^1^ Diseases were categorized using International Classification of Diseases (ICD) 10 code, and only the primary ICD10 discharge code was consistently available for each hospitalization event. Data from Brazil were selected in this study because it has one of the most comprehensively-collected hospitalization databases in the world and has high geographic resolutions. Brazil also has interesting geographic characteristics, capturing a broad range of income/development levels and climatic zones. There are 27 states and 5 regions in the country, which largely reflect climatic differences, as well as sub-national variations in human development, as based on income, longevity, and education. For the senior age group, two states in the North region were dropped from analysis, because ICD10 chapters included in the model had fewer than 10 cases/month on average in the study period. Therefore, 27 states were included in analysis for the young age group, while the old age group had 25 states. Preprocessing of the hospitalization data, such as an adjustment for the rapid shift in the number of hospitalizations due to the change in coding practice in 2008, was done as described in our previous study.^2^

### eAppendix 2: Additional information on down-sampled data

To assess how the number of events in a time series may affect predictions from various quantitative models, we performed a down sampling analysis.^3,4^ From the national-level data, we randomly subsampled the time series of ICD10 chapters, including all-cause pneumonia hospitalizations, using binomial distribution with various rates such that

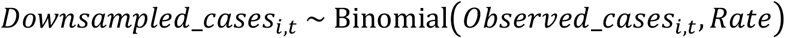

where *Observed_cases_i,t_* was the actual number of hospitalizations for disease *i* reported in time period *t* at the national-level. Cases were randomly sampled at rates of 10%, 1%, and 0.25%, to simulate the population sizes of different regions and states in Brazil. For example, the population sizes of the five regions were between 5% and 48% of the entire national population of infants and seniors. At the state level, the smallest state represented only 0.3% (State 14 in the North region; n=10,500) for children under 12 months of age and 0.2% (State 12 in the North region; n=5,900) of people 80+ years of age across Brazil.

We repeated this sampling process 100 times and created 100 down-sampled datasets at each of these rates. We then fit the models to each of these down-sampled datasets and quantified the impact of the vaccine. To evaluate the effect of sample size on the performance of the models, the estimated impact of the vaccine from the down-sampled data were compared to those from the national-level data, assuming the national estimate was the “ground truth.”

### eAppendix 3: Additional information on the simulated time series data

The number of outcome (*Outcome_t_*) and four control diseases (*ControlA_t_, ControlB_t_, ControlC_t_*, and *ControlD_t_*) are randomly sampled from the following means to generate the time series:

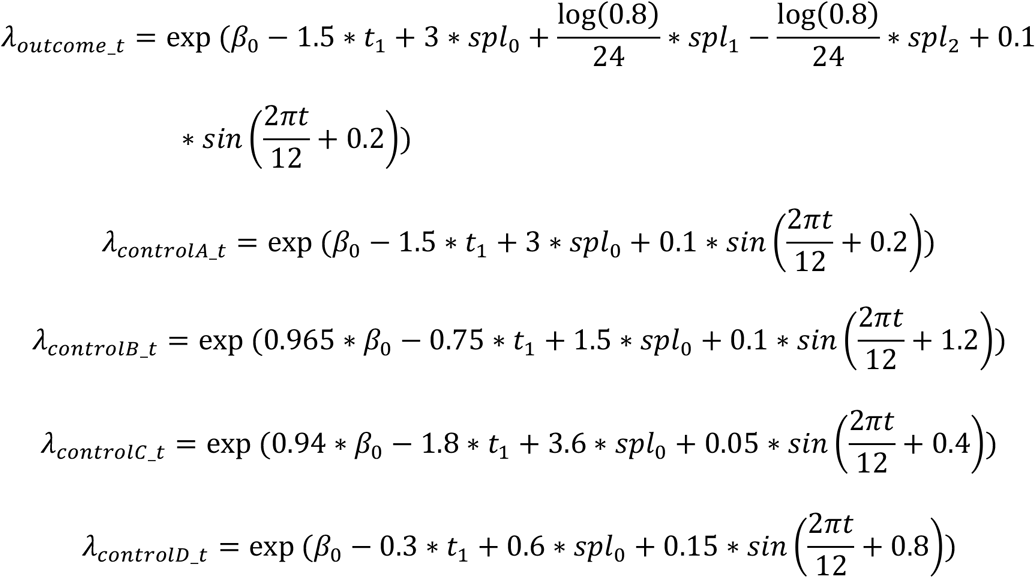

*spl*_0_ captures a secular trend that begins in month 43 (specified as a linear spline). *spl*_1_ and *spl*_2_ capture a vaccine-associated decline, which continues for 24 months and then levels out (also specified as linear splines).

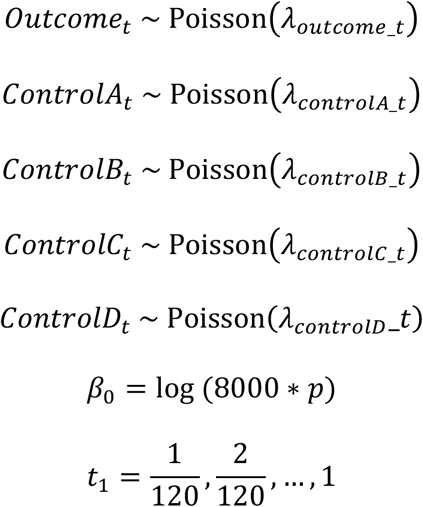

where *p* is the sample size (1, 0.1, 0.01, or 0.0025). For each value of *p*, we simulated 100 time series data. *ControlA_t_* is considered “perfect control,” as it had the exact same trend as the outcome until the simulated vaccine introduction.

### eAppendix 4: Additional information on the synthetic control model

The SC modeling framework is as follows:

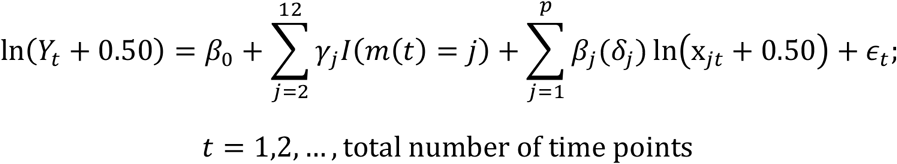

where *Y_t_* represents the number of pneumonia hospitalizations at time *t*; x_*jt*_ represents the count of control disease *j* at time *t*; *m*(*t*) is a function that maps a time point to the corresponding calendar month; *γ_j_* represents the month *j* regression coefficient; *I*(.) represents the indicator function; *p* is the total number of control diseases in the analysis; *β_j_*(*δ_j_*) is the regression coefficient for control disease *j* which is given a spike-and-slab prior distribution (depending on *δ_j_*) in order to allow for data-driven variable selection^5^; *δ_j_* are binary random variables that indicate if control disease *j* is included in the regression model (*δ_j_* = 1) or not (*δ_j_* = 0); and 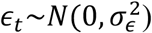. The regression coefficients, *β_j_*(*δ_j_*) for control disease *j*, are not time varying, because we assume that the relationships between the outcome and control diseases are constant over time. Time series for the outcome and control diseases were log-transformed prior to being used in the model in order to alleviate the effects of epidemics on the long-term trend and to more closely resemble normality assumed in the model. As a continuity correction, 0.5 was added to all data points. The 2009 influenza pandemic was adjusted for by including dummy variables for the months in which the pandemic peaked.^1^

Full details on the prior distribution can be found in the paper by Bruhn, *et al*.^1,5^ Briefly, we used spike and slab priors to select variables in the candidate models with equal prior probability of inclusion for each variable (π = 0.5). We collected 9,000 posterior samples after a burn-in period of 1,000 iterations for each fitted model. The convergence of the Markov chain Monte Carlo sampling algorithm was evaluated using the Geweke diagnostic and Gelman Rubin diagnostic.^6–8^ There were no obvious signs of non-convergence as the p-values from the Geweke diagnostic tests were all larger than 0.05 and the potential scale reduction factors were <1.1 across all parameters. The effective sample size was also checked to confirm that we had enough posterior samples to make valid inference.^9,10^

### eAppendix 5: Additional information on the STL+PCA model

The seasonal-trend decomposition procedure based on locally weighted scatterplot smoothing (STL method) decomposes time series into three components: trend, seasonality, and the remaining variation in data (Figure 1).^11^ The observed number of control disease *j* cases in month *t*, denoted by *ControlDisease_jt_*, can be written as follows:

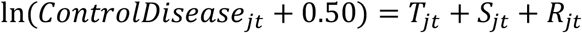

where *T_jt_*, *S_jt_*, and *R_jt_* are the trend component, annual seasonal component, and remainder component, respectively. For the STL trend extraction, the span of the locally weighted scatterplot smoothing (LOESS) window can be changed to control the smoothness of the extracted trends. The larger the LOESS window is, the smoother the extracted trends are (eFigure 4). We set it to be 5, 25, and 59 months, and selected the optimal size using the deviance information criterion (DIC).^12^ The model with the optimal window size was then used to generate the counterfactual for the post-vaccine period and to quantify the impact of vaccine.

The principal component analysis (PCA) allows us to create new uncorrelated projections that explain the maximum variability in the data overall.^13–16^ PCA was applied to the trends for control diseases extracted by the STL method. Extracted trends were first converted into standard deviation units (Z scores) by subtracting off the mean of each variable and dividing by its standard deviation, as the variables included in PCA need to have the same scale. PCA involved an *n*×*p* matrix, *X*, where *p* represents the number of control diseases and *n* represented the total number of time points. The *j*th column of matrix *X, x_j_*, represents the time series of a scaled extracted trend for control disease *j*. We then find a linear combination of these extracted trends (i.e., columns of matrix *X*) with maximum variance, which are given by

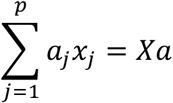

where *a* is a vector of constants *a*_1_, *a*_2_, …, *a_p_*. The variance of this linear combination can be written as follows:

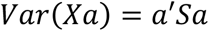

where *S* represents the sample covariance matrix associated with the dataset. Therefore, the linear combination that explains the largest variance can be identified by obtaining a *p*-dimensional vector *a* which maximizes the quadratic form *a′Sa*.

The JAGS model was fit using the rjags package in R version 3.4.3 (Vienna, Austria).^17^ We initialized two independent Markov chains and collected 10,000 posterior samples after a burn-in period of 5,000 iterations for the Brazil data, and 5,000 posterior samples after a burn-in period of 1,000 iterations for the simulated time series data. Convergence was assessed as described in eAppendix 4. The JAGS model was specified as follows. A negative binomial regression was used due to the over-dispersion present in the data:

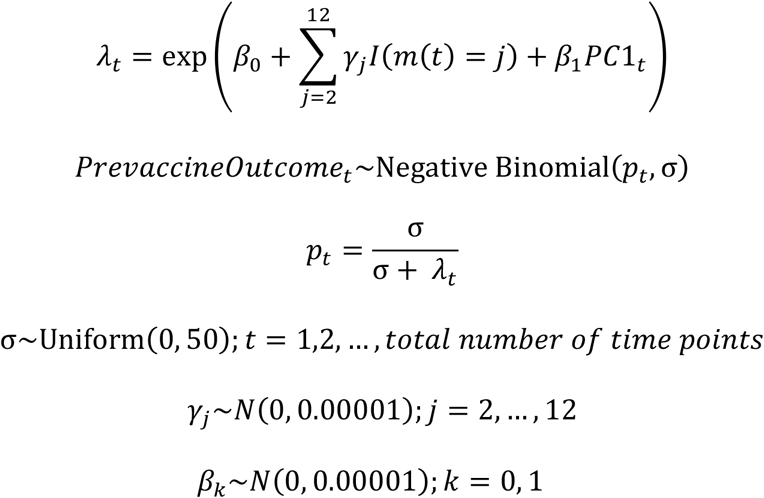

where *PrevaccineOutcome_t_* represents the number of pneumonia hospitalizations at time *t* in the pre-vaccine period; p_t_ is a probability parameter specific to time period *t*; *m*(*t*) is a function that maps a time point to the corresponding calendar month; *γ_j_* represents the month *j* regression coefficient; *I*(.) represents the indicator function; *β*_0_ is an intercept; *β*_1_ is the regression coefficient for the first principal component (PC1); σ is a dispersion parameter for the negative binomial model; *λ_t_* is the expected value of the negative binomial distribution; and 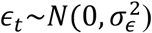.

### eAppendix 6: Time-trend model

The framework for the time-trend model is as follows:

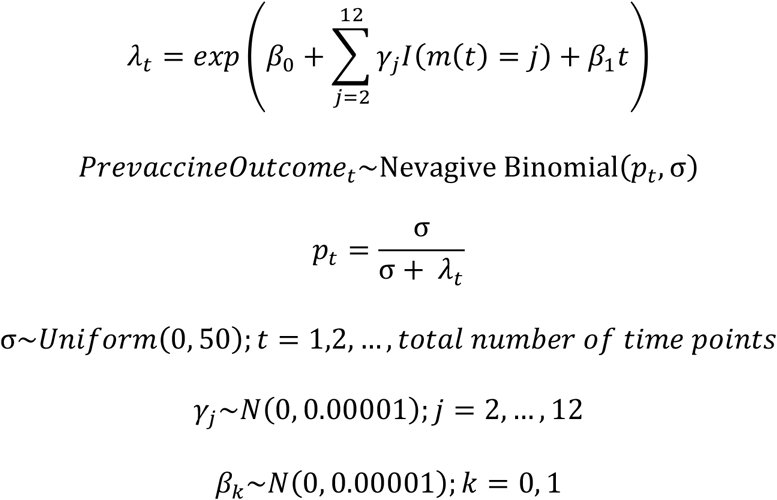

The definitions of the parameters are the same as the STL+PCA model (eAppendix 5).

The JAGS model was fit using the rjags package in R version 3.4.3 (Vienna, Austria). ^7^ We initialized two independent Markov chains and collected 10,000 posterior samples after a burn-in period of 5,000 iterations for the Brazil data, and 5,000 posterior samples after a burn-in period of 1,000 iterations for the simulation data. Convergence was assessed as described in eAppendix 4. The model was fit to the pre-vaccine data and extended to the post-vaccine period to generate counterfactual.

### eAppendix 7: Mean squared error

The performance of models was compared using the mean squared error (MSE), which was calculated as follows:

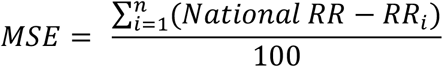

where *RR_i_* represents a rate ratio for dataset *i* and *n* is the number of total datasets compared in each analysis (i.e., total number of states for state-level analysis and 100 for down-sampled analysis).

### eAppendix 8: Results on cross validation of the STL+PCA model

We have performed cross validation using the down-sampled pre-vaccine data with the rate 0.25% for the elderly in Brazil. We trained our model using four years of pre-vaccine data (2004–2007) and generated prediction for 2008. Although it was part of the pre-vaccine period, we avoided using the 2009 data due to a large 2009 influenza outbreak. The 95% credible interval of the rate ratio included one in 97% of 100 down-sampled datasets, suggesting that this model successfully predicted the 2008 data.

## Notes

**Description of conflicts of interest:** D.M.W. has previously received an investigator-initiated research grant from Pfizer and consulting fees from Pfizer, Merck, GSK, and Affinivax. L.S. and R.J.T. have an ownership interest in Sage Analytica, a research consultancy with government, nongovernment, and pharmaceutical industry clients, including an investigator-initiated research grant from Pfizer (completed in 2013). Other authors do not have conflict of interest.

**Sources of financial support:** This study was supported by the Bill & Melinda Gates Foundation Grant OPP1114733. K.S. was also supported by the Funai Foundation for Information Technology through the Funai Overseas Scholarship. D.M.W. was also supported by NIH/National Institute on Aging Grants P30AG021342 (Scholar at the Claude D. Pepper Older Americans Independence Center at Yale University School of Medicine), R01AI123208, and R56AI110449 (NIH/NIAID); and NIH National Center for Advancing Translational Science Grants CTSA UL1TR001863.

